# Altered actin filament dynamics in the *Drosophila* mushroom bodies lead to fast acquisition of alcohol consumption preference

**DOI:** 10.1101/623991

**Authors:** Andrew R. Butts, Shamsideen A. Ojelade, Alexandra Seguin, Collin B. Merrill, Aylin R. Rodan, Adrian Rothenfluh

## Abstract

Alcohol use is highly prevalent in the United States and across the world, and every year millions of people suffer from alcohol use disorders (AUDs). While the genetic contribution to developing AUDs is estimated to be 50-60%, many of the underlying molecular mechanisms remain unclear. Previous studies from our lab revealed that *Drosophila* lacking RhoGAP18B and Ras Suppressor 1 (Rsu1) display reduced sensitivity to ethanol-induced sedation. Both Rsu1 and RhoGAP18B are negative regulators of the small Rho-family GTPase, Rac1, a modulator of actin dynamics. Here we investigate the role of Rac1 and its downstream target, the actin-severing protein cofilin, in alcohol consumption preference. We show that these two regulators of actin dynamics can alter experience-dependent alcohol preference in a bidirectional manner: expressing either activated Rac1 or dominant-negative cofilin in the mushroom bodies (MB) abolishes experience-dependent alcohol preference. Conversely, dominant-negative Rac1 or activated cofilin MB expression lead to faster acquisition of alcohol preference. Our data show that Rac1 and cofilin activity are key to determining the rate of acquisition of alcohol preference, revealing a critical role of actin dynamics regulation in the development of voluntary self-administration in *Drosophila*.

**Significance Statement:** The risks for developing an alcohol use disorder (AUD) are strongly determined by genetic factors. Understanding the genes and molecular mechanisms that contribute to that risk is therefore a necessary first step for the development of targeted therapeutic intervention. Here we show that regulators of actin cytoskeleton dynamics can bidirectionally determine the acquisition rate of alcohol self-administration, highlighting this process as a key mechanism contributing to the risk of AUD development.

## Introduction

Alcohol use is highly prevalent; in the United States, 56% of adults reported they were monthly drinkers, and in 2015, 15 million adults were diagnosed with alcohol use disorder (AUD). The misuse of alcohol results in 88,000 annual deaths and results in a $249 billion economic burden (NSDUH, 2015; CDC-ARDI, 2006-2010; Sacks et al., 2015). Up to 60% of the risk of developing AUD can be attributed to genetic predisposition (Dick et al., 2006; Gelernter and Kranzler, 2011), and numerous genes are associated with human AUD phenotypes in genome-wide association studies. However, the genes from these association studies often do not group into obvious over-represented categories of physiological function (e.g. Evangelou et al., 2018), leaving many of the molecular pathways of AUD obscure.

Numerous model organisms have been established to investigate these genetic mechanisms. *Drosophila melanogaster*, the vinegar fly, is one genetically tractable model organism that has been used to model addiction-like behaviors (Rodan and Rothenfluh, 2010; Kaun et al., 2012). Flies naïvely respond to alcohol similarly to humans: low doses result in loss of inhibition and hyperactivity, while high doses result in loss of postural control and sedation (Wolf et al., 2002; Lee et al., 2008). Flies will also learn to prefer alcohol and voluntarily choose to consume it over a period of 5 days (Devineni and Heberlein, 2009). Initially, flies display naïve aversion to ethanol, but a pre-exposure to vaporized alcohol leads to preferential ethanol self-administration (Peru y Colon de Portugal et al., 2014). Numerous genes whose mutations cause altered self-administration in *Drosophila* also have human orthologs with polymorphisms that associate with AUD phenotypes (Grotewiel and Bettinger, 2015; Ojelade et al., 2015a; Gonzales et al., 2018).

One of the pathways linked to behavioral responses to alcohol is the dynamic regulation of the actin cytoskeleton, which centers around the Rho-family of small GTPases (Rothenfluh and Cowan, 2013). Loss of RhoGAP18B in flies, a negative regulator of Rac1 GTPase, leads to changes in ethanol-induced sedation or hyperactivity, depending on the RhoGAP18B isoform affected (Rothenfluh et al., 2006). Loss of a second negative regulator of Rac1, Rsu1, also causes altered sensitivity to ethanol-induced sedation, as well as altered alcohol self-administration in a 5-day preference assay (Ojelade et al., 2015a). Here, we investigate the role of Rac1, and its downstream mediator, cofilin – an actin-severing protein – in experience-dependent alcohol preference (EDAP). Using a behavioral paradigm that separates an alcohol pre-exposure from the consummatory choice, we show that proper regulation of actin-dynamics is required during the acquisition for preference to develop. In a new preference paradigm that allows close temporal observation of consummatory behavior, we then show that genetic manipulation that causes the opposite effect on actin dynamics leads to an accelerated acquisition of alcohol consumption preference. These bidirectional phenotypes emphasize the critical role regulators of actin-dynamics play in voluntary alcohol self-administration.

## Materials and Methods

### Fly Stocks and Genetics

Male flies were used for all experiments. Flies were grown and kept on standard cornmeal/agar medium at 25°C with 70% relative humidity. Flies were outcrossed to the *w* Berlin* genetic background for at least 5 generations, with the exception of *MB-GeneSwitch*, for which sibling-matched controls were used to equalize their genetic background in our behavioral assays. Transgenic flies were obtained from the Bloomington Stock Center: *UAS-Rac*^*CA*^ (stock BL# 6291), *UAS-Rac*^*DN*^ (BL# 6290), *UAS-tsr*^*CA*^ (BL# 9236), *UAS-tsr*^*DN*^ (BL# 9238). *MB-GeneSwitch* and *MB247-Gal4* were gifts from Dr. Gregg Roman.

### Drug Feeding

Manipulating whole-fly actin dynamics was achieved via feeding the flies either jasplakinolide (JPK) or latrunculin A (Lat.A). JPK and Lat.A were dissolved in 100% DMSO. Flies were food deprived for 16 hr before feeding on 250 mM liquid sucrose with either 200 nM JPK, 18.9 µM Lat.A, or equivalent volume of DMSO vehicle. For the *MB-GeneSwitch* experiments, flies were 16 hr food deprived before feeding on 250 mM liquid sucrose with or without 0.5 mM mifepristone (aka RU-486). 50 mM mifepristone was dissolved in 95% ethanol and diluted for feeding, diluting the ethanol to a sub-preference inducing alcohol concentration for feeding.

### Ethanol Pre-Exposures

Groups of 10 males were exposed to ethanol vapor and air via the Booze-O-Mat assay described previously (Wolf et al., 2002). 20 min exposures occurred 24 hr prior to CAFÉ assays. Control flies were mock-exposed to air vapor without ethanol.

### 16-hr CAFÉ Assay

Ethanol preference was performed via the 2-bottle choice Capillary Feeder (CAFÉ) assay as described (Devineni and Heberlein 2009) with modifications. Our CAFÉ assay consisted of a 6-well plate with 4 small holes drilled for the insertion of pipette tips and 20 µl capillaries (VWR, Radnor, PA). Capillaries were filled via capillary action, and a small mineral oil overlay was added to reduce evaporation. Preference assays with 10 males per well were conducted at 25° C and 70% relative humidity and flies chose between liquid sucrose food with or without 15% ethanol. For the *MB-GeneSwitch* experiments, food-deprived flies were fed 0.5 mM mifepristone for 3 hr prior to the CAFÉ assay.

### 16-hr COLA Assay

The COLA (for CAFÉ-based Online Learning Assay) apparatus was based on the above described CAFÉ assay, but 4-well plates were used with 2 capillaries per well. The solutions offered were 250 mM sucrose with or without 15% ethanol. COLA assays were recorded with a time-lapse camera set to a 5 min interval (TLC200, Brinno Inc., Taipei City, Taiwan). Video recordings were then binned into 2 hr intervals to calculate preference for each interval.

### 30-min FRAPPE Assay

Naïve alcohol aversion was tested in a 30 min 2-choice preference assay called the FRAPPE (for Fluorometric Reading Assay of Preference Primed by Ethanol). This assay was performed as previously described (Peru y Colon de Portugal et. al., 2014). Groups of 35 male flies chose between 340 mM liquid sucrose food with or without 15% ethanol, after a 6 hr food deprivation.

### G/F-actin In Vivo Assay

G/F-actin assay was performed according to the manufacturer’s instructions (G/F-actin In Vivo Assay Kit, Cytoskeleton, Denver, CO). G-and F-actin bands on western blots were scanned by densitometry and the ratios of free G-actin to F-actin were calculated.

### Experimental Design and Statistical Tests

Analysis of the experiments was performed using Prism 8 (GraphPad Software Inc., San Diego, CA). The data were tested for normality by examining the QQ plots and using the Wilks-Shapiro normality test, which showed that the data were normally distributed with the exception of Figure 5. Data from the FRAPPE experiments were not normally distributed, as we previously found (Peru y Colon de Portugal et al., 2014), therefore these data were analyzed with non-parametric statistics. Data with an *n* greater than 8 were checked for outliers, defined as >2.5 standard deviations outside the mean for single point measures (in Figures 1-4, 5 out of 575 data points were excluded), and 1.5x the interquartile range outside the upper and lower quartiles for nonparametric data (Figure 5: 3 of 192 data points excluded). For repeat measures, we excluded runs where more than half the time points lay >2 standard deviations outside the mean (Figure 6: 1 of 47 runs excluded).

**Figure 1.**
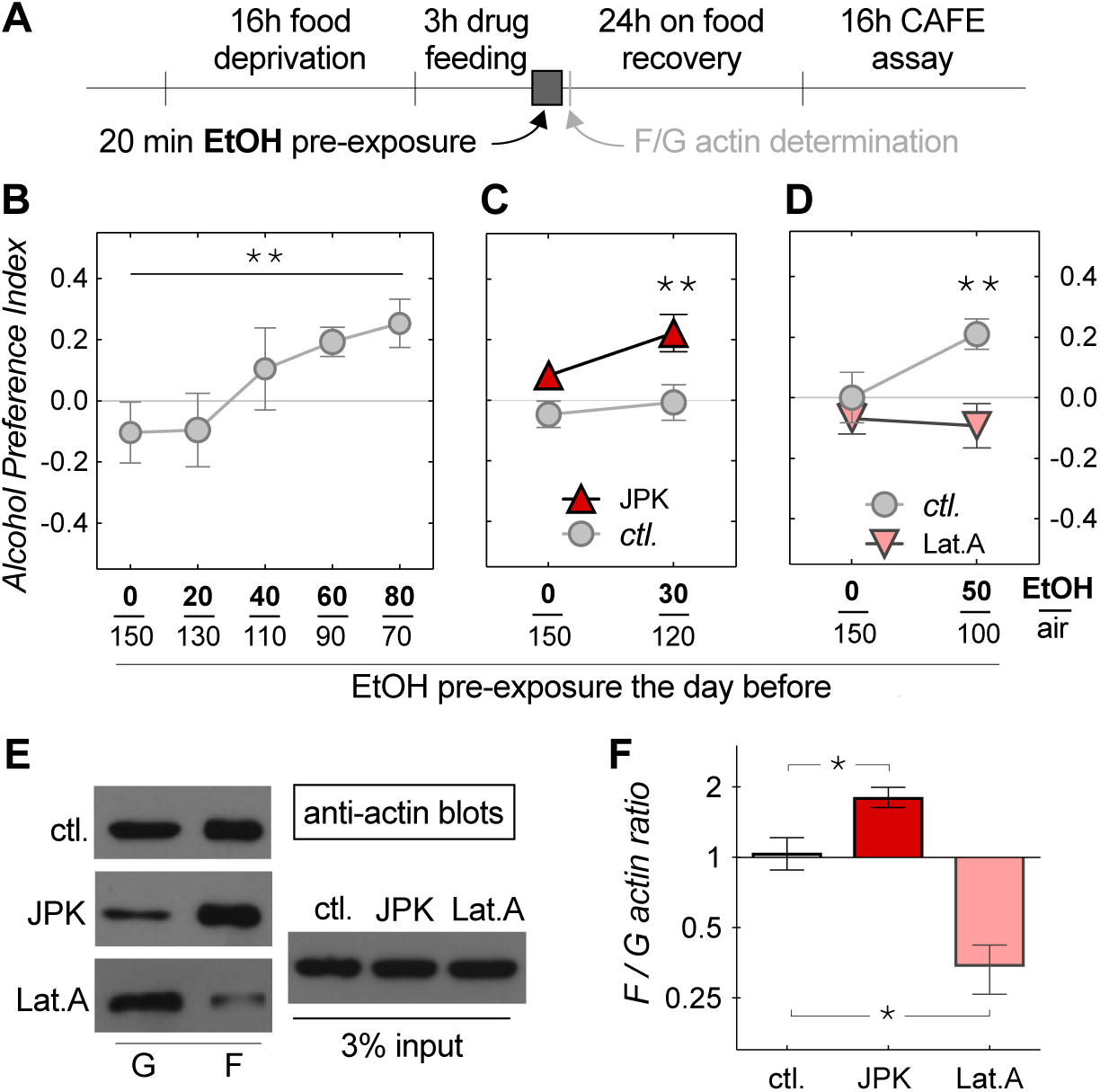
Effects of altered actin polymerization on experience-dependent alcohol preference (EDAP). ***A***, Schematic of the experimental design. Flies were food deprived for 16 hr prior to a 3-hr feeding on control food or food with either 200 nM jasplakinolide (JPK) to increase, or with 8.0 ug/ml latrunculin A (Lat.A) to decrease the F/G-actin ratio. Following the drug feeding, flies were pre-exposed to an ethanol vapor and air mixture for 20 min, and then allowed to recover for 24 hr on standard fly food. Flies were then assayed in a 16-hr abbreviated CAFÉ assay. ***B***, Control flies show a dose-dependent switch from alcohol avoidance to alcohol preference (slope of linear regression 0.005±0.003 95% CI, significantly different from 0, \*\**p* = 0.0028, *n* = 6 per dose). Ethanol vapor to air ratios for the pre-exposures are displayed on the X-axis. ***C***, Pre-feeding on JPK caused EDAP at a sub-threshold pre-exposure dose (two-way ANOVA, with a significant effect, *F*(1,44) = 11.9, *p* = 0.0012 of JPK. Sidak’s multiple post-hoc comparison \*\**p* = 0.006 at 30/120 EtOH/air, *n* = 12 per data point). ***D***, Pre-feeding on Lat.A prevented EDAP at a dose which induced preference in control flies (two-way ANOVA, with a significant effect, *F*(1,44) = 7.9, *p* = 0.0074 of Lat.A. Sidak’s multiple post-hoc comparison \*\**p* = 0.0046 at 50/100 EtOH/air, *n* = 12). ***E***, Anti-actin western-blots run from whole fly extracts. Blots on left show globular (G) and filamentous (F) actin fractions. The input extract shown on the right. ***F***, Western blot quantification shows significant, predicted changes in F/G-actin ratios (one-way ANOVA, *F*(2,6) = 24.5, *p* = 0.0013, with Dunnett’s post-hoc multiple comparisons, *p < 0.027, *n* = 3; shown in this, and following Figures – except Fig.5 – are means with standard errors).

**Figure 2.**
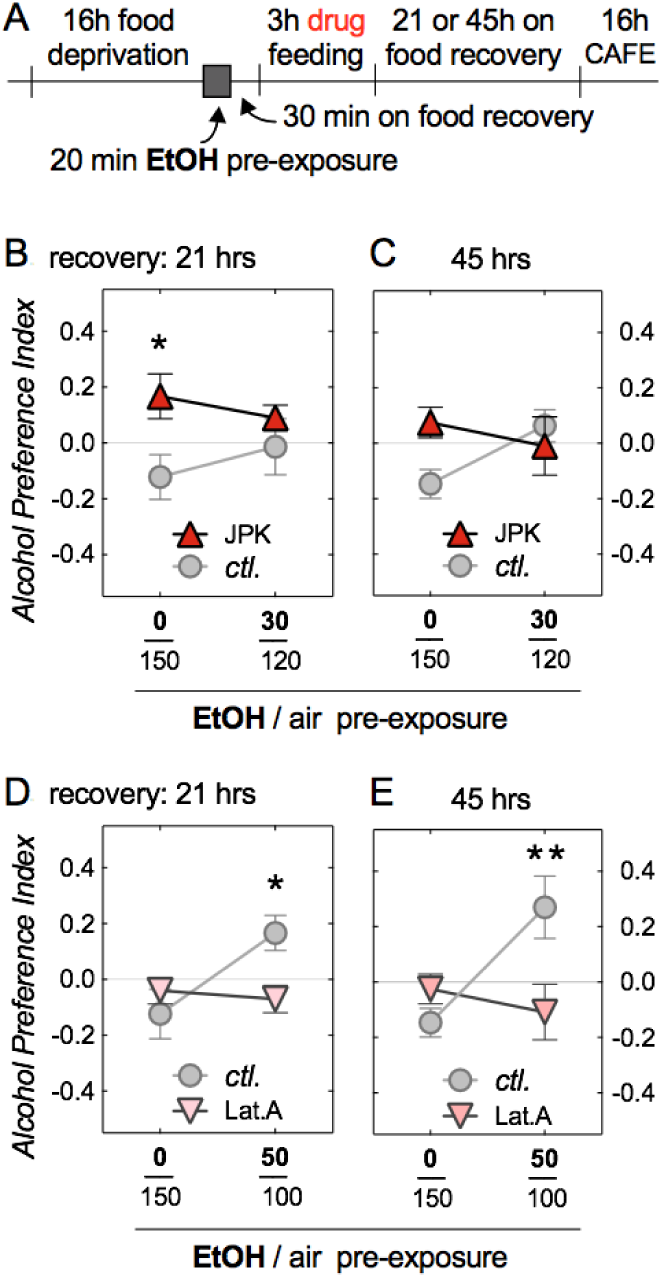
Effects of actin polymerization post-acquisition on EDAP. ***A***, Experimental design similar to Fig.1 with the exception that flies were drug-fed after alcohol vapor pre-exposure, and that two recovery times were tested. ***B*** and ***C***, Feeding JPK after alcohol exposure led to naive preference in mock ethanol-exposed flies 21 hr (two-way ANOVA, with a significant effect, *F*(1,32) = 4.8, *p* = 0.037, of JPK. Sidak’s multiple post-hoc comparison \**p* = 0.018 at 0/150 EtOH/air, *n* = 6,12 per data point), but not 45 hr post-feeding. Ethanol-exposed flies showed no effect of JPK at either recover time. ***D*** and ***E***, Feeding Lat.A after alcohol exposure prevented EDAP in ethanol-exposed flies 21 hr (two-way ANOVA, with a significant interaction, *F*(1,20) = 2.3, *p* = 0.021. Sidak’s multiple post-hoc comparison \**p* = 0.033 at 50/100 EtOH/air, *n* = 6), and 45 hr post-feeding (two-way ANOVA, with a significant interaction, *F*(1,20) = 8.8, *p* = 0.0075. Sidak’s multiple post-hoc comparison \*\**p* = 0.0094 at 50/100 EtOH/air, *n* = 6).

**Figure 3.**
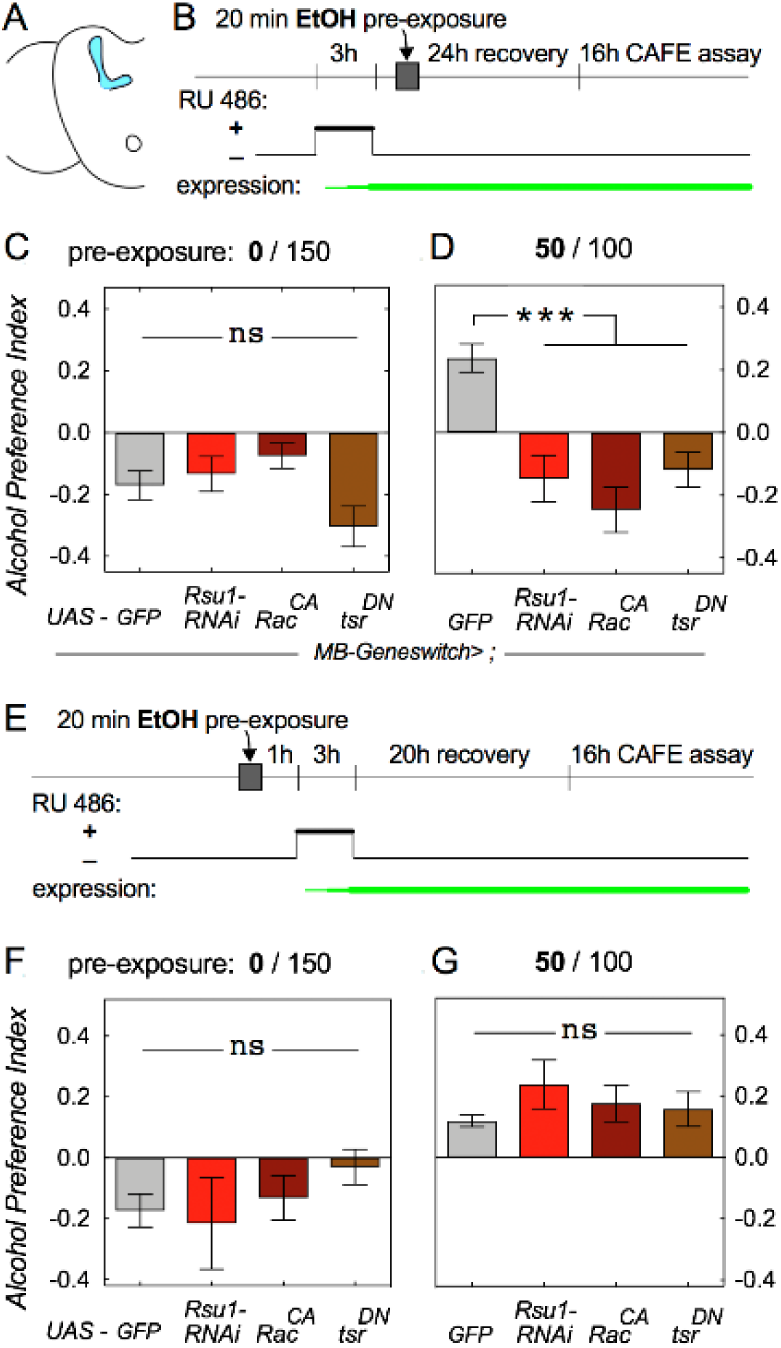
Decreasing F-actin turnover in the mushroom bodies (MB) during pre-exposure suppresses EDAP. ***A***, Brain schematic highlighting the MB. ***B***, Experimental design with RU486 feeding prior to ethanol exposure to activate the *MB-GeneSwitch* Gal4 driver. ***C***, Decreasing F-actin turnover via three different transgenes did not alter naïve alcohol avoidance (one-way ANOVA, *F*(3,56) = 2.6, *p* = 0.06, *n* = 12, or 24 for GFP control). ***D***, Decreasing F-actin turnover with transgene activation prior to ethanol pre-exposure prevented EDAP (one-way ANOVA, *F*(3,73) = 15.0, *p* < 0.0001; Sidak’s multiple post-hoc comparison \*\*\**p* ≤ 0.0008, *n* = 12-18, and 29 for GFP control). ***E***-***G***, Activating *MB-GeneSwitch* after alcohol pre-exposure had no effect on naïve avoidance (one-way ANOVA, *F*(3,32) = 1.9, *p* = 0.15, *n* = 6, or 18 for GFP control in *F*), or on EDAP (one-way ANOVA, *F*(3,32) = 0.94, *p* = 0.44, *n* = 6, or 17 for GFP control in *G*).

**Figure 4.**
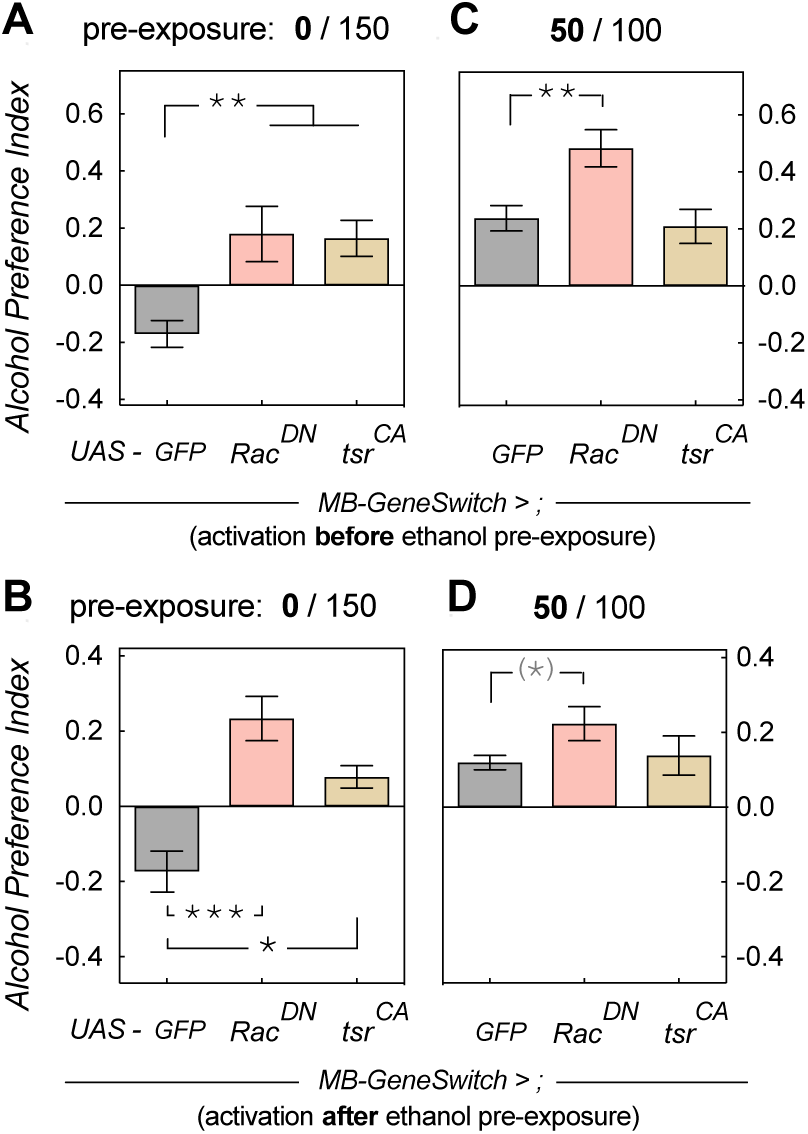
Increasing F-actin turnover in the MB leads to naïve alcohol preference. ***A*** and ***B***, Increasing F-actin turnover with two different transgenes led to naïve preference in mock-exposed flies irrespective of *MB-GeneSwitch* activation before (one-way ANOVA, *F*(2,45) = 10.6, *p* = 0.0002; Sidak’s multiple post-hoc comparison \*\**p* = 0.0007, *n* = 12, or 24 for GFP control in *A*) or after alcohol pre-exposure (one-way ANOVA, *F*(2,27) = 11.2, *p* = 0.0003; Sidak’s multiple post-hoc comparison \*\*\**p* = 0.0003, \**p* = 0.021, *n* = 6, or 18 for GFP control in *B*). ***C***, Increasing F-actin turnover prior to ethanol pre-exposure in alcohol pre-exposed flies still led to EDAP, and in the case of Rac^DN^, enhanced preference (one-way ANOVA, *F*(2,56) = 6.7, *p* = 0.0025; Sidak’s multiple post-hoc comparison \*\**p* = 0.0031, *n* = 12,18 or 29 for GFP control). ***D***, Activating *MB-GeneSwitch* after pre-exposure also still allowed development of EDAP. One-way ANOVA indicated no differences (*F*(2,27) = 2.8, *p* = 0.075), while a t-test with Bonferroni correction suggested significant enhancement of EDAP with Rac^DN^ (*t* = 2.5, *df* = 24, ^(*)^*p* = 0.0426, effect size: Hedge’s *g* = 0.95, 0.11-1.80 95% CI, *n* = 9,17, and 6 for *tsr*).

**Figure 5.**
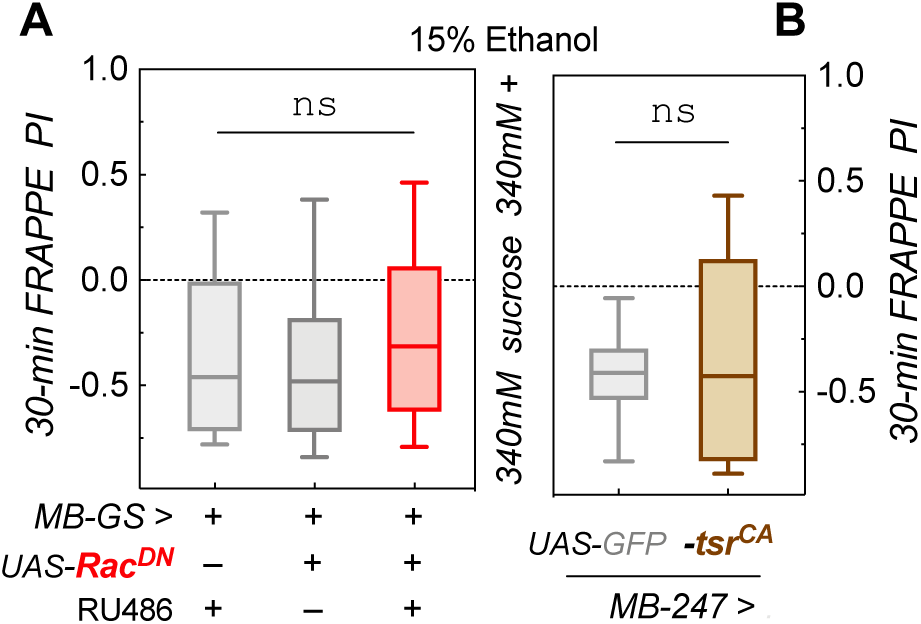
Increased MB F-actin turnover does not affect naïve alcohol avoidance in a short preference assay. ***A***, Adult expression of *UAS-Rac*^*DN*^ with *MB-GeneSwitch* did not alter avoidance in a 30-min preference assay (Kruskal-Wallis test statistic = 2.09, *p* = 0.35, *n* = 47 or 48; shown are medians with quartile boxes and 10-90% whiskers). ***B***, Constitutive *UAS-tsr*^*CA*^ expression with *MB247-Gal4* did not change naïve alcohol avoidance either (Mann-Whitney test, *U* = 216, *p* = 0.86, *n* = 14, 32).

**Figure 6.**
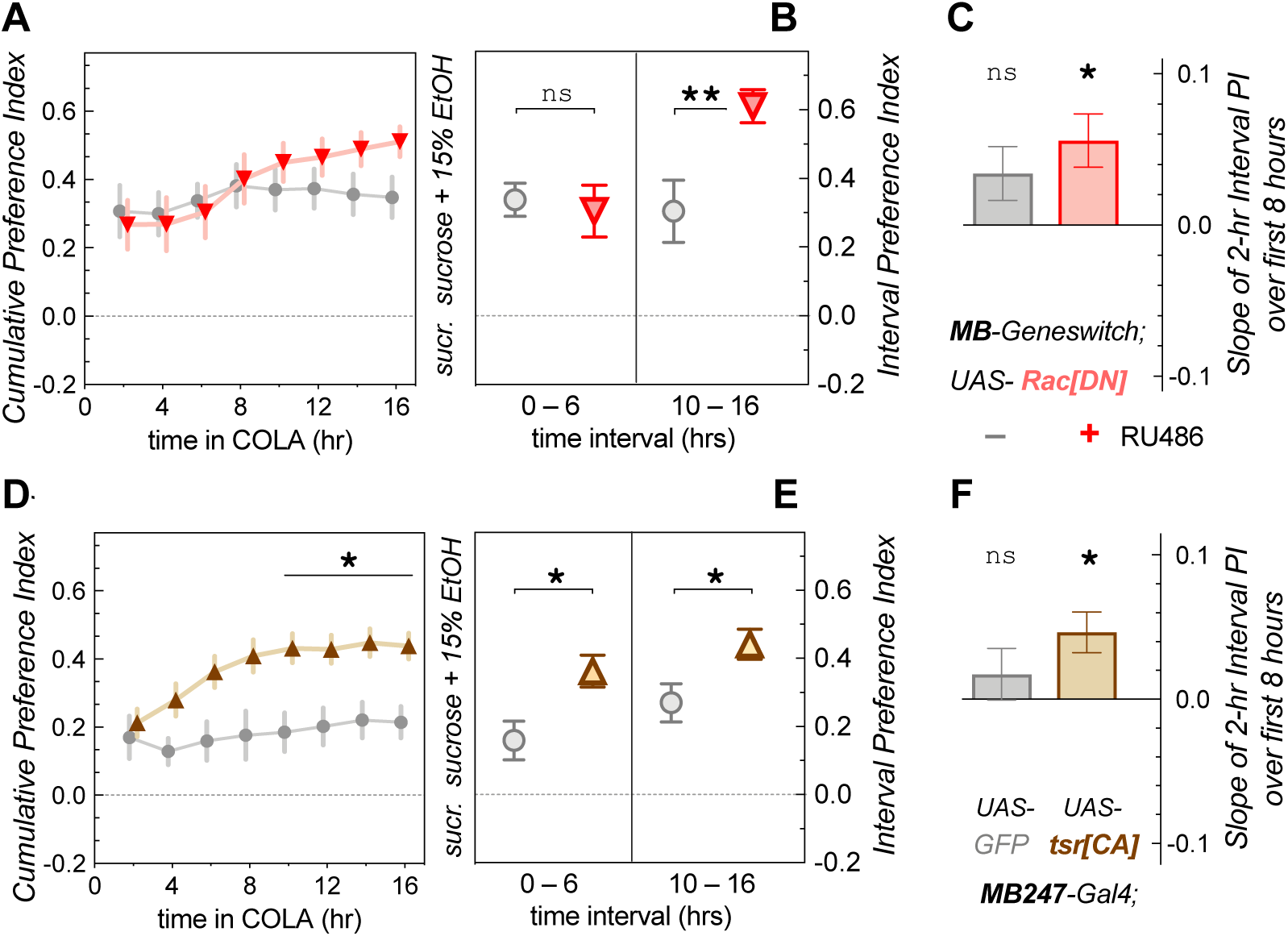
Increasing F-actin turnover in the MB causes fast acquisition of alcohol preference. Experiments were conducted in a CAFÉ online learning assay which monitored consumption from the food capillaries with 5-min resolution. ***A***, Cumulative preference index, from time 0 to time X, for adult-expressed *UAS-Rac*^*DN*^ in the MB showed a significant effect of time (two-way RM ANOVA for repeat measures, *F*(2.118, 42.37) = 7.6, *p* = 0.0012) and of interaction (time × RU486 treatment, *F*(7,140) = 3.1, *p* = 0.0042, n = 11), but no post-hoc differences (Sidak’s multiple comparisons, *t* < 2.3, *p* > 0.27). ***B***, Interval preference index for the first and last six hours revealed no difference at the start (one-way ANOVA with 2 pre-selected Sidak’s pairwise comparisons, *t* = 0.35, *p* = 0.93), but a significant increase for the treated group (*t* = 3.17, *p* = 0.0058). ***C***, Average slope of the linear regression for the 2-hr interval preferences in the first 8 hours. The RU486-treated group showed a positive average slope (one sample t-test with Bonferroni threshold adjustment, *t* = 1.91, ^ns^*p* = 0.085 for – group, *t* = 3.18, \**p* = 0.0098 for + group). ***D***, Cumulative preference index when expressing *MB247>tsr*^*CA*^ showed significant effects of time (two-way RM ANOVA, F(1.180, 40.70) = 7.8, p = 0.0017), treatment (*UAS-transgene, F*(1,22) = 10.4, *p* = 0.0039) and interaction (*F*(7,154) = 2.95, *p* = 0.0062). The preference at numerous time points was significantly increased with *UAS-tsr*^*CA*^ (Sidak’s multiple comparisons, *t* > 3.3, \**p* < 0.027). ***E***, Interval preference indices were also increased during both the first six hours (one-way ANOVA with 2 pre-selected Sidak’s pairwise comparisons, *t* = 2.81, \**p* = 0148) and the last six hours (*t* = 2.37, \**p* = 0.0436). ***F***, The average slope of the linear regression in the first 8 hours was positive for the experimental *tsr*^*CA*^, but not GFP control group (one sample t-test with Bonferroni threshold adjustment, *t* = 0.97, ^ns^*p* = 0.35 for GFP group, *t* = 3.31, \**p* = 0.0069 for *tsr*^*CA*^ group).

## Results

### F-actin polymerization is involved in experience dependent alcohol preference

Our previous studies showed that wild-type flies naively avoid alcohol and choose to consume a sucrose solution over a sucrose solution containing alcohol in a 2-choice paradigm (Peru y Colon de Portugal et al., 2014). This alcohol avoidance changes to preferential alcohol consumption over multiple days (Devineni and Heberlein, 2009) or with a pre-exposure to alcohol vapor the day before (Peru y Colon de Portugal et al., 2014). Here we recapitulate that result, showing that wild-type flies show slight naive avoidance to alcohol, but this avoidance switches to preference in a dose-dependent manner with the level of alcohol pre-exposure 24-hr prior to the start of a 16-hr abbreviated CAFÉ assay (Fig. 1*B*, slope of linear regression 0.005±0.003 95% CI, significantly different from 0, \*\**p* = 0.0028, *n* = 6 per dose). Studies from our lab showed that manipulating regulators of actin dynamics, including Rac1 GTPase, can lead to behaviorally distinct alcohol phenotypes, including alcohol-induced sedation and locomotion activation (Rothenfluh et al., 2006; Ojelade et al., 2015a; Ojelade et al., 2015b). Here, we wanted to investigate whether the manipulation of the state of F-actin would affect preferential alcohol consumption. Pre-feeding wild-type flies 200 nM jasplakinolide (JPK), a peptide with actin polymerization activity (Fig. 1*E* and *F*, one-way ANOVA, *F*(2,6) = 24.5, *p* = 0.0013, with Dunnett’s post-hoc multiple comparisons, \**p* < 0.027, *n* = 3), was able to facilitate experience-dependent alcohol preference (EDAP) at a sub-threshold pre-exposure dose of ethanol that did not induce preference in control flies (Fig. 1*C*, two-way ANOVA, with a significant effect, *F*(1,44) = 11.9, *p* = 0.0012 of JPK. Sidak’s multiple post-hoc comparison \*\**p* = 0.006 at 30/120 EtOH/air, *n* = 12 per data point). Conversely, pre-feeding flies 18.9 µM latrunculin A (Lat.A), a toxin that disrupts actin filaments and increases G-actin (Fig. 1*E* and *F*), prevented the development of EDAP after alcohol pre-exposure at a higher dose (Fig. 1*D*, two-way ANOVA, with a significant effect, *F*(1,44) = 7.9, *p* = 0.0074 of Lat.A. Sidak’s multiple post-hoc comparison \*\**p* = 0.0046 at 50/100 EtOH/air, *n* = 12). These experiments show that feeding flies drugs interfering with F-actin polymerization prior to alcohol vapor pre-exposure alters their alcohol preference.

We next wanted to investigate whether proper regulation of actin dynamics is required during the 20-min alcohol pre-exposure, or during the 16-hr preference test for normal EDAP development. Feeding flies Lat.A after the alcohol pre-exposure still prevented EDAP when tested 21, or 45 hr after the drug feeding (Fig. 2*D*, two-way ANOVA, with a significant interaction, *F*(1,20) = 2.3, *p* = 0.021. Sidak’s multiple post-hoc comparison \**p* = 0.033 at 50/100 EtOH/air, *n* = 6 and Fig. 2*E*, two-way ANOVA, with a significant interaction, *F*(1,20) = 8.8, *p* = 0.0075. Sidak’s multiple post-hoc comparison \*\**p* = 0.0094 at 50/100 EtOH/air, *n* = 6). The effects of JPK, however, were more complicated: JPK-feeding no longer facilitated EDAP upon sub-threshold alcohol pre-exposure, except for one time point, where it resulted in naïve alcohol preference without alcohol pre-exposure (Fig. 2*B*, two-way ANOVA, with a significant effect, *F*(1,32) = 4.8, *p* = 0.037, of JPK. Sidak’s multiple post-hoc comparison \**p* = 0.018 at 0/150 EtOH/air, *n* = 6,12 per data point and Fig. 2*C*). These data confirmed the importance of proper actin dynamics for EDAP, but its requirement during or after the pre-exposure was harder to interpret given the mixed results we obtained with JPK vs. Lat.A. Furthermore, our feeding experiments were systemic, therefore complicating interpretation in light of pleiotropic alcohol phenotypes that can be obtained depending on which neurons are affected (Rothenfluh et al., 2006; Ojelade et al., 2015a).

### Increased mushroom body F-actin stability during pre-exposure suppresses EDAP

Previously, we have shown that Ras suppressor 1 (Rsu1) is a negative regulator of the small GTPase Rac1. Rsu1 suppresses GTP loading of Rac1 and leads to an increase in F-actin (Ojelade et al., 2015a) via inactivation of the actin-severing protein cofilin (Ojelade et al., 2015b). We used three manipulations known to increase F-actin: *UAS-Rsu1-RNAi* to knock down *Rsu1, UAS-Rac*^*CA*^, to overexpress activated Rac1, and *UAS-tsr*^*DN*^ to overexpress dominant-negative, inactive cofilin. We previously showed that Rsu1 and Rac1 are required in the mushroom bodies (MB), a known center for associative learning (Cognigni et al., 2018), to develop alcohol preference in a 4-day CAFÉ assay (Ojelade et al., 2015a). We therefore restricted the genetic manipulations to the MB using the drug-inducible Gal4, *MB-GeneSwitch (MB-GS)*, which is activated upon consumption of mifepristone (aka RU486, Fig. 3*A* and *B*). First, we activated *MB-GS* prior to alcohol pre-exposure, which increases F-actin prior to acquisition of an alcohol preference (Fig. 3*B*). As expected, naïve flies avoided alcohol in the 16-hr CAFÉ (Fig. 3*C*). Following exposure to a preference-inducing dose of alcohol, F-actin mutants failed to develop preference, while the control, *MB-GS>UAS-GFP*, developed EDAP (Fig. 3*D*, one-way ANOVA, *F*(3,73) = 15.0, *p* < 0.0001; Sidak’s multiple post-hoc comparison \*\*\**p* ≤ 0.0008, *n* = 12-18, and 29 for GFP control), consistent with our previous findings (Ojelade et al., 2015a). To determine whether reduced F-actin dynamics, and increased F-actin, had a role after the acquisition of preference, we activated *MB-GS* after the alcohol pre-exposure (Fig. 3*E*). As before, naïve flies avoided alcohol in the 16hr CAFÉ (Fig. 3*F*). In contrast, all the F-actin mutants and control flies developed preference after alcohol pre-exposure (Fig. 3*G*, one-way ANOVA, *F*(3,32) = 0.94, *p* = 0.44, *n* = 6, or 17 for GFP control). Taken together, these data indicate proper actin dynamics is required during alcohol pre-exposure for the acquisition of EDAP.

### F-Actin turnover in the mushroom bodies produces naïve preference for alcohol

We next investigated the effect of increased F-actin turnover on EDAP. To that end we expressed two transgenes in the MB using *MB-GS*. The first was a dominant negative mutation of Rac1, which keeps Rac1 bound to GDP and inactive. The second manipulation was a constitutively active mutation in *tsr* producing an activated cofilin protein. As before, these manipulations were restricted to the MB using *MB-GS*. Activating expression of *MB-GS* before, or after alcohol pre-exposure did not disrupt EDAP, and in the case of Rac^DN^ even caused an increase in preference (Fig. 4*C*, one-way ANOVA, *F*(2,56) = 6.7, *p* = 0.0025; Sidak’s multiple post-hoc comparison \*\**p* = 0.0031, *n* = 12,18 or 29 for GFP control and Fig. 4*D*, One-way ANOVA indicated no differences *F*(2,27) = 2.8, *p* = 0.075). Surprisingly, overexpression of these mutants also caused preference in naive flies, regardless of whether we induced *MB-GS* activation before (Fig. 4*A*, one-way ANOVA, *F*(2,45) = 10.6, *p* = 0.0002; Sidak’s multiple post-hoc comparison \*\**p* = 0.0007, *n* = 12, or 24 for GFP control) or after alcohol exposure (Fig. 4*B*, one-way ANOVA, *F*(2,27) = 11.2, *p* = 0.0003; Sidak’s multiple post-hoc comparison \*\*\**p* = 0.0003, \**p* = 0.021, *n* = 6, or 18 for GFP control). The transgenes’ effects were very strong, with significant effect sizes (Hedge’s *g*) between 1.2 and 1.8 (Fig. 4*A* and *B*). This was unexpected for two reasons. First our results with the opposite mutations (Fig. 3) suggested that actin dynamics during, not after, the pre-exposure are critical for preference development. Second, and even more surprising, our prior findings suggested that the MB are critical for EDAP, but that neurons outside the MB are involved in mediating and determining naïve alcohol avoidance (Ojelade et al., 2015a). This led us to two alternative hypotheses, the first that increased F-actin turnover in the MB does in fact affect naïve alcohol avoidance. The second hypothesis was that enhanced F-actin turnover gradually increases the voluntary consumption of alcohol compared to controls specifically during our 16-hr ‘testing’ period.

### Increased F-Actin turnover mutants display normal naïve alcohol aversion

In order to investigate our first hypothesis, we assayed our increased F-actin turnover mutants in a short 30-min 2-choice preference assay, called FRAPPE. This assay pairs food offerings with a fluorometric dye, which can be quantified through the cuticle of the fly abdomen to determine which food was preferred (Peru y Colon de Portugal et al., 2014). Flies with adult-induced expression of *MB-GS>UAS-RAC*^*DN*^ showed no difference in naïve alcohol aversion compared to control flies (Fig. 5*A*, Kruskal-Wallis test statistic = 2.09, *p* = 0.35, *n* = 47 or 48; shown are medians with quartile boxes and 10-90% whiskers). Since the timing of expression of the mutants had no impact on the phenotype in Figure 4, we also used a constitutive MB-specific driver, *MB247-Gal4*. As seen with Rac^DN^, *MB247-Gal4>UAS-tsr*^*CA*^ flies also displayed naïve alcohol aversion in our 30-min preference assay, as did the control flies (Fig. 5*B*, Mann-Whitney test, *U* = 216, *p* = 0.86, *n* = 14, 32). Therefore, reduced naïve alcohol aversion did not explain why these mutants showed preference in our 16-hr CAFÉ assay in the absence of prior ethanol exposure.

### Increased F-actin turnover in the MB accelerates the acquisition of alcohol preference

We designed our 16-hr CAFÉ assay as the preference testing phase of our EDAP paradigm. However, our above results suggested that mutants with increased F-actin turnover might start out avoiding alcohol, but then acquire preference within this 16-hr ‘testing’ period. In order to investigate this idea, we had to develop an assay that could monitor preference in real time, as opposed to checking consumption and preference at the end of the 16 hour “testing’ period, or every 24 hours as for long-term CAFÉ assays (Ja et al., 2007; Devineni and Heberlein, 2009). We therefore developed the CAFÉ-based online learning assay (COLA), which differs from a traditional CAFÉ because consumption of the food offerings is video recorded in real time, allowing minute-level resolution of feeding behavior. Because feeding in the COLA is sporadic, and not continuous, we decided to check preference and consumption every 2 hr. Flies with induced *MB-GS>Rac*^*DN*^ showed a gradual increase of the preference index (PI) over the 16-hr experiment, which was less evident for the uninduced control (Fig. 6*A*, two-way RM ANOVA for repeat measures, *F*(2.118, 42.37) = 7.6, \*\**p* = 0.0012 significant effect of time and of interaction, time × RU486 treatment, *F*(7,140) = 3.1, \*\**p* = 0.0042, *n* = 11). When we calculated an interval PI for the first and last six hours of the experiment, experimental Rac^DN^ flies showed a significantly higher PI at the end of the experiment than controls, showing that they acquired a preference increase for the duration of the 16-hr experiment (Fig. 6*B*, one-way ANOVA with 2 pre-selected Sidak’s pairwise comparisons, *t* = 3.17, \*\**p* = 0.0058). Because the PI change was most obvious in the first 8 hours, we also determined the slope of a linear regression curve through the first 4 2-hour PI intervals for each replicate, essentially asking whether there is an increase in alcohol preference within the first 8 hours of the experiment. The average slope of these linear regressions was significantly positive for the induced experimental Rac^DN^ flies, but not for the uninduced controls (Fig. 6*C*, one sample t-test with Bonferroni threshold adjustment, *t* = 1.91, ^ns^*p* = 0.085 for – group, *t* = 3.18, \**p* = 0.0098 for + group). It is noteworthy that both control and experimental flies showed a fairly high, albeit indistinguishable, preference even within the first 2 hours of the experiment. There are two reasons for that: first, flies show more alcohol aversion in our 30-min assay (Fig. 5) which offers the food in numerous wells, and not small capillaries like in the CAFÉ. This is partially due to their aversion of the (relatively undiluted) smell of alcohol (data not shown). The other reason was the genetic background of the *MB-GS* driver, which showed relatively high initial preference in the COLA setup, irrespective of RU486 induction or *UAS-transgene* presence. We therefore switched to the *MB247-Gal4* driver to test the effect of activated cofilin mutants. Experimental *MB247>tsr*^*CA*^ again showed increased preference over the 16-hr experiment compared to controls (Fig. 6*D*, two-way RM ANOVA, F(1.180, 40.70) = 7.8, \*\**p* = 0.0017 for time; and *F*(1,22) = 10.4, \*\**p* = 0.0039 for genotype; and (*F*(7,154) = 2.95, \*\**p* = 0.0062 for the genotype × time interaction). The preference at numerous time points was significantly increased with *UAS-tsr*^*CA*^ (Fig. 6*D*, Sidak’s multiple comparisons, *t* > 3.3, \**p* < 0.027), and their PI was significantly higher than the control PI during both the first (Fig. 6*E*, one-way ANOVA with 2 pre-selected Sidak’s pairwise comparisons, *t* = 2.81, \**p* = 0.0148) and the last six hours of the assay (*t* = 2.37, \**p* = 0.0436). Lastly, experimental flies’ average slope for the linearly-regressed first 4 2-hr interval PIs was also positive while the control’s was not (Fig. 6*F*, one sample t-test with Bonferroni threshold adjustment, *t* = 0.97, ^ns^*p* = 0.35 for GFP group, *t* = 3.31, \**p* = 0.0069 for *tsr*^*CA*^ group), indicating that *MB247>tsr*^*CA*^ flies learn to prefer alcohol within the first 8 hours of the assay (also visually evident in Fig. 6*D*).

## Discussion

### Requirement for mushroom body F-actin turnover in the development of alcohol preference

Proper dynamic regulation of the actin cytoskeleton is known to be relevant for learning and memory (Lamprecht, 2016), including the development of drug preference (Rothenfluh and Cowan 2013). Our systemic drug feeding experiments in Fig.1 suggested that increasing F-actin facilitates the development of EDAP, while decreasing F-actin abolished preference. However, when we restricted the manipulations of F-actin to the MB, our data suggested that increased MB F-actin abolished EDAP, while decreased F-actin enhanced EDAP. This discrepancy may be explained by two possibilities. First, the MB are a well-known center for learning and memory in flies (Cognigni et al., 2018) and have been shown to be involved in alcohol consumption preference (Xu et al., 2012; Kaun et al., 2011). Conversely, we have shown that neurons outside the MB can also affect alcohol preference (Ojelade et al., 2015a). Therefore, altering F-actin systemically changes actin dynamics both in MB and other neurons, and the resulting behavioral output is the sum of the different circuits affecting behavioral responses. Thus changes in the MB exclusively might lead to a different phenotype than systemic changes. An alternative explanation would be the actual mechanism of action of Lat.A compared to cofilin, which both lead to decreased F-actin. Lat.A binds to G-actin monomers and prevents actual polymerization of actin filaments (Morton et al., 2000), whereas cofilin causes severing of actin filaments, but does not prevent new formation of filaments. Indeed, cofilin activity increases actin treadmilling and membrane protrusion (Kanellos and Frame, 2016). Thus, Lat.A may be viewed as decreasing overall dynamics of F-actin turnover, whereas activated cofilin increases F-actin turnover and re-generation. The relevant change induced by these manipulations may not be in the overall F/G-actin ratio, but rather the dynamic ability for the F-actin cytoskeleton to change on demand. Indeed, the mechanism of action for JPK is not to just rigidly stabilize actin filaments, but also to enhance the rate of F-actin nucleation (Bubb et al., 2000). Together, these results suggest that the proper development of EDAP requires enhanced F-actin dynamics and turnover in the *Drosophila* MB, which is counteracted by Rac^CA^ (Fig.3*D*). Similarly, Dietz and colleagues (2012) found that overexpression of Rac^CA^ in the rodent nucleus accumbens reduced cocaine-induced place preference, highlighting the importance of this GTPase in drug abuse across phyla.

### The temporal requirement of proper F-actin dynamics in preference development

Our behavioral paradigm consists of an alcohol pre-exposure followed by a preference test. We were therefore able to ask whether proper F-actin dynamics is required during the pre-exposure and acquisition of preference, or at later stages, i.e. testing, or consolidation. Our data with reduced F-actin turnover (Fig.3) suggested that enhanced F-actin dynamics is required during the acquisition of preference, but not thereafter. However, decreasing F-actin with systemic Lat.A feeding after the pre-exposure also led to prevented EDAP (Fig.2*D* and *E*). Thus, some F-actin generation, or at the very least, F-actin stabilization may also be required during the consolidation or recall phase of alcohol-induced consumption preference. Indeed, injection of Lat.A into the rodent basolateral amygdala after training and consolidation of methamphetamine-induced conditioned place preference (CPP) abolished the expression/recall of that place memory (Young et al., 2014), whereas injection of Lat.A into the nucleus accumbens shell led to reduced consolidation of morphine-induced CPP (Li et al., 2015). These data suggest that proper F-actin dynamics is important during numerous stages of addiction-related behavior, including during the acquisition of drug-preference memory, as our data suggests.

### Fast acquisition of preferential alcohol self-administration with enhanced F-actin turnover

Enhancing F-actin dynamics led to facilitated alcohol preference, even in the absence of an alcohol pre-exposure (Fig.4*A* and *B*). Our subsequent analysis showed that this is not due to altered naive avoidance of alcohol (Fig.5; Peru y Colon de Portugal et al., 2014), but rather because these flies showed fast acquisition of alcohol preference. We determined this using our novel COLA assay, which allows for much refined temporal resolution of monitoring self-administration. The parsimonious explanation for the fast acquisition would be that increased F-actin dynamics enhances the acquisition of the alcohol preference. Supporting this interpretation is the finding that overexpression of dominant-negative Rac or activated cofilin in the nucleus accumbens was able to induce CPP with a sub-threshold dose of cocaine (Dietz et al., 2012), suggesting that these molecular manipulations facilitate the acquisition of a drug-induced memory.

A number of reports have suggested that activated Rac leads to active forgetting of shock-conditioned odor avoidance, while dominant-negative Rac and activated cofilin reduce the rate of forgetting (Shuai et al., 2010; Cervantes-Sandoval et al., 2016). Our results could be interpreted in a similar light, with *MB>Rac*^*CA*^ flies not displaying EDAP (Fig.3*D*), because they forgot it right away. Conversely, *MB>Rac*^*DN*^ and *tsr*^*CA*^ showed enhanced preference (Fig.4,6), because they did not forget any of it. However, induction of Rac^CA^ after ethanol pre-exposure did not affect EDAP (Fig.3*E*), showing that activated Rac does not drive an active reduction/forgetting of the acquired ethanol-memory. Rac^DN^ affected shock-induced odor memory up to 24 hours after the memory induction (Shuai et al, 2010), thus it seems unlikely that the timing of *Rac*^*CA*^ expression after the ethanol pre-exposure is the cause for the lack of a *Rac*^*CA*^ effect on alcohol preference. The observed difference to shock-conditioned odor avoidance may simply be a reflection of the different memories, one stemming from aversive punishment, and the other from appetitive reinforcement. Even within appetitive conditioning, numerous reports suggest that drug-induced memories are stronger, or even categorically different from sucrose-induced ones. These drug-memories are unaffected by manipulations affecting F-actin that do affect food memories (Kiraly et al., 2010; Young et al., 2014; Laguesse et al., 2017). Regardless of the exact function of this pathway leading to activated cofilin and F-actin dynamics in suppressing forgetting and/or facilitating memory acquisition, we here show that this pathway can bidirectionally affect the development of alcohol self-administration preference. Our data also demonstrate that increased F-actin turnover in the *Drosophila* MB can facilitate the acquisition of drug memories, suggesting a high degree of evolutionary conservation of this pathway’s function in the development of substance abuse disorders.

## Acknowledgements

The authors thank lab members for continued discussion of experiments and suggestions on the manuscript. Supported by NIH T32DA007290, F31AA021340 (S.A.O.), AHA 16CSA28530002, NIH R01DK110358 (A.R.R.), NIH R01AA019526 (A.R.), U2M2 (A.R.R., A.R.) and the University of Utah Neuroscience Initiative (A.R.).

